# Robust Neural Networks are More Interpretable for Genomics

**DOI:** 10.1101/657437

**Authors:** Peter K. Koo, Sharon Qian, Gal Kaplun, Verena Volf, Dimitris Kalimeris

## Abstract

Deep neural networks (DNNs) have been applied to a variety of regulatory genomics tasks. For interpretability, attribution methods are employed to provide importance scores for each nucleotide in a given sequence. However, even with state-of-the-art DNNs, there is no guarantee that these methods can recover interpretable, biological representations. Here we perform systematic experiments on synthetic genomic data to raise awareness of this issue. We find that deeper networks have better generalization performance, but attribution methods recover less interpretable representations. Then, we show training methods promoting robustness – including regularization, injecting random noise into the data, and adversarial training – significantly improve interpretability of DNNs, especially for smaller datasets.

## 1. Introduction

As powerful function approximators that autonomously learn features, deep neural networks (DNNs) have been applied to learn genomic sequence patterns that are predictive of a regulatory function, such as protein binding, chromatin accessibility, and histone marks (Zhou & Troyanskaya, 2015; Quang & Xie, 2016; Kelley et al., 2016; Hiranuma et al., 2017; Alipanahi et al., 2015; Koo et al., 2018). To interpret a trained DNN, attribution methods – which include *in silico* mutagenesis (Alipanahi et al., 2015; Zhou & Troyanskaya, 2015), backpropagation to the inputs (Simonyan et al., 2013), Deeplift (Shrikumar et al., 2016), SHAP (Lundberg & Lee, 2017), guided backprop (Springenberg et al., 2014), and integrated gradients (Sundararajan et al., 2017) – provide importance scores (to first order approximation) for individual nucleotides (nts).

One factor that tends not to be considered is the quality of the DNN’s fit beyond test performance, *i.e*. the smoothness of the decision boundary. Deeper networks are more expressive (Raghu et al., 2016), which allows them to fit more complicated functions, but also enables them to easily overfit to “noisier” functions. Nevertheless, DNNs that learn noisier functions to attain perfect accuracy on the training set may still generalize well, irrespective of whether they are regularized (Zhang et al., 2016).

It has also been observed that DNNs are susceptible to adversarial perturbations (Goodfellow et al., 2014). This observation was made in the context of image recognition, where a small – often imperceptible to the human eye – change to an input image leads to a drastically different classification by the model. Adversarial examples can be generated using iterative gradient-based methods to perturb the natural data. To defend against these adversarial examples, an effective method is adversarial training, where adversarial examples are computed at each epoch and injected into the dataset during training of the neural network (Madry et al., 2017). Many methods to generate adversarial examples have been proposed (Dong et al., 2017; Moosavi-Dezfooli et al., 2016; Goodfellow et al., 2014). These methods have been shown to be effective in increasing the *robustness* of DNNs.

Motivated by this line of work, we apply similar ideas in the context of regulatory genomic datasets and explore how interpretability improves using methods aimed to promote robustness, including regularization, random noise injection, and adversarial training. We perform systematic experiments on synthetic DNA sequences to test the efficacy of a DNN’s ability to learn combinations of sequence motifs that comprise so-called regulatory codes. We find that reliability of gradient-based attribution methods varies significantly with the depth of the network, even though the classification performance is similar. We also find that training procedures that promote robustness have a small impact on classification performance, but can significantly improve the interpretability of the model.

## 2. Experimental overview

We posit that robustness may not necessarily affect generalization performance, but is indicative of interpretability with gradient-based attribution methods. To test this, we created a synthetic dataset that recapitulates a simple regulatory code classification task.

### Dataset

We generated 30,000 synthetic sequences by embedding known motifs in specific combinations. Positive class sequences were synthesized by embedding 3 to 5 “core motifs” – randomly selected with replacement from a pool of 10 position frequency matrices, which include the forward and reverse complement motifs for CEBPB, Gabpa, MAX, SP1, and YY1 (Mathelier et al., 2016) – along a random sequence model. Negative class sequences were generated following the same steps with the exception that the pool of motifs include 100 non-overlapping “background motifs” from the JASPAR database (Mathelier et al., 2016). Background sequences can thus contain core motifs; however, it is unlikely to randomly draw motifs that resemble a positive regulatory code. We randomly combined synthetic sequences of the positive and negative class and randomly split the dataset into training, validation and test sets with a 0.7, 0.1, and 0.2 split, respectively. Availability of dataset and code: github.com/p-koo/uncovering_regulatory_codes

### Models

Leveraging recent progress on representation learning of genomic sequence motifs (Koo & Eddy, 2018), we designed two convolutional neural networks (CNNs), namely LocalNet and DistNet, to learn “local” representations (whole motifs) and “distributed” representations (partial motifs), respectively. Both take as input a 1-dimensional one-hot-encoded sequence with 4 channels, one for each nt (A, C, G, T), and have a fully-connected (dense) output layer with a single sigmoid activation. The hidden layers for each model are:

1. LocalNet
  1. convolution (24 filters, size 19, stride 1, ReLU) max-pooling (size 50, stride 50)
  2. fully-connected layer (96 units, ReLU)
2. DistNet:
  1. convolution (24 filters, size 7, stride 1, ReLU)
  2. convolution (32 filters, size 9, stride 1, ReLU) max-pooling (size 3, stride 3)
  3. convolution (48 filters, size 6, stride 1, ReLU) max-pooling (size 4, stride 4)
  4. convolution (64 filters, size 4, stride 1, ReLU) max-pooling (size 3, stride 3)
  5. fully-connected layer (96 units, ReLU)

We created two variations of each model, without regularization and with regularization. For the regularized models, we incorporate batch normalization (Ioffe & Szegedy, 2015) in each hidden layer; dropout (Srivastava et al., 2014) with probabilities corresponding to: LocalNet (layer1 0.1, layer2 0.5) and DistNet (layer1 0.1, layer2 0.2, layer3 0.3, layer4 0.4, layer5 0.5); and *L*2-regularization on all parameters in the network with a strength equal to 1e-6.

### Training

We uniformly trained each model by minimizing the binary cross-entropy loss function with mini-batch stochastic gradient descent (100 sequences) for 100 epochs. We updated the parameters with Adam using default settings (Kingma & Ba, 2014). All reported performance metrics are drawn from the test set using the model parameters which yielded the lowest loss on the validation set.

### Attribution methods

To test interpretability of trained models, we generate attribution scores by employing backprop from the logits – prior to the sigmoid activation – to the inputs (Simonyan et al., 2013). We also employ smoothgrad (Smilkov et al., 2017), a technique that builds upon standard backprop to mitigate noise in attribution scores. Smoothgrad adds Gaussian noise to the inputs and then averages the resulting attribution scores. In practice, we generate 50 noisy samples for each sequence by adding noise drawn from a Gaussian distribution 𝒩 (0, 0.1) to each nt variant. We note that while the inputs are no longer categorical, we do not expect pathological behavior from the model since DNNs treat the inputs as continuous values.

### Quantifying interpretability

Since we have the ground truth of embedded motif locations in each sequence, we can test the efficacy of attribution scores. To quantify the interpretability of a given attribution map, we calculate the area under the receiver-operator characteristic curve (AU-ROC) and the area under the precision-recall curve (AU-PR), comparing the distribution of attribution scores where ground truth motifs have been implanted (positive class) and the distribution of attribution scores at positions not associated with any ground truth motifs (negative class). Specifically, we first multiply the attribution scores (*S*_*ij*_) and the input (*X*_*ij*_) and reduce the dimensions to get one score per position (see Fig. 1A), according to *C*_*j*_ = ∑_*i*_ *S*_*ij*_*X*_*ij*_, where *i* is the alphabet and *j* is the position. We then calculate the information of the sequence model, *M*_*ij*_, according to *I*_*j*_ = log_2_ 4 ∑_*i*_ *M*_*ij*_ log_2_ *M*_*ij*_. Positions that are given a positive label are defined by *I*_*j*_ > 0, while negative labels are given by *I*_*j*_ = 0. The AU-ROC and AU-PR is then calculated separately for each sequence using the distribution of *C*_*j*_ at positive label positions against negative label positions. For reference, Figure 1B shows representative examples of attribution maps and ground truth for various AU-ROC and AU-PR values.

**Figure 1.**
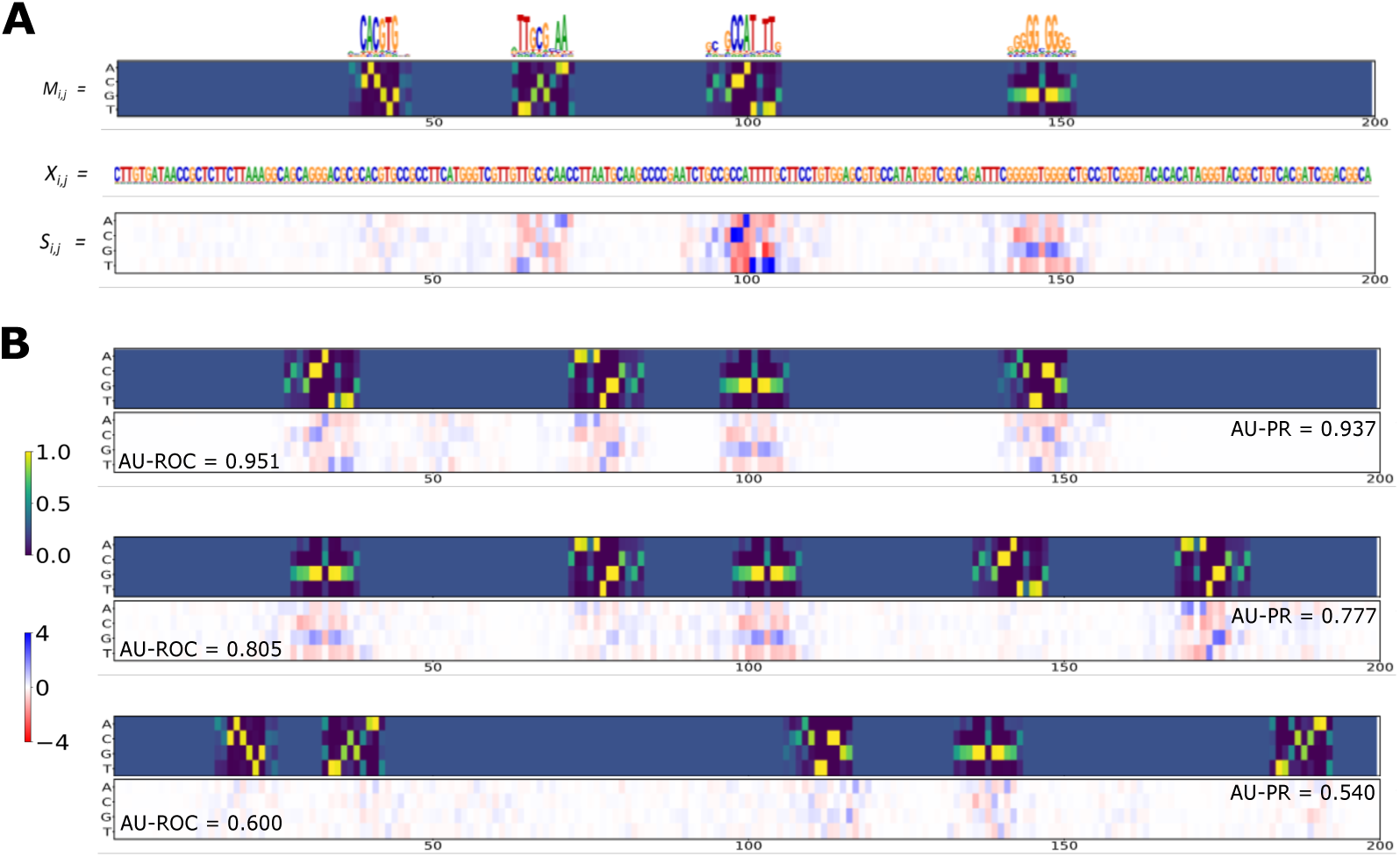
Quantifying interpretability. (A) Shows a representative sequence model *M*, the sequence generated from the model (*X*), and an attribution map from a trained DNN (*S*). (B) Various examples of attribution maps by the same model (LocalNet with regularization) for different interpretability scores given by AU-ROC (bottom left inset) and AU-PR (top right inset). The ground truth sequence model is shown above for visual comparison.

## 3. Results

We trained LocalNet and DistNet with and without regularization and compared the classification performance using the area the receiver-operator characteristic curve (AUC). We also compared the interpretability performance using the AU-ROC and AU-PR of attribution score distributions using ground truth from sequence models.

### Accuracy does not translate to interpretability

Classification AUC is comparable between DistNet (0.964) and LocalNet (0.967). However, LocalNet is significantly more interpretable with a higher AU-ROC (0.751± 0.055) and AU-PR 0.599±0.112) compared to DistNet which yields 0.561± 0.068 and 0.442± 0.105, respectively. We surmise that DistNet, which has a higher expressivity to fit more complicated functions (Raghu et al., 2016), has fit to a “noisier” function, resulting in poorer interpretability with gradient-based attribution methods. Smoothgrad is designed to address this issue by sampling the gradients about the local function of the input data; however, this technique seems to only marginally improve interpretability (Table 1).

**Table 1.**
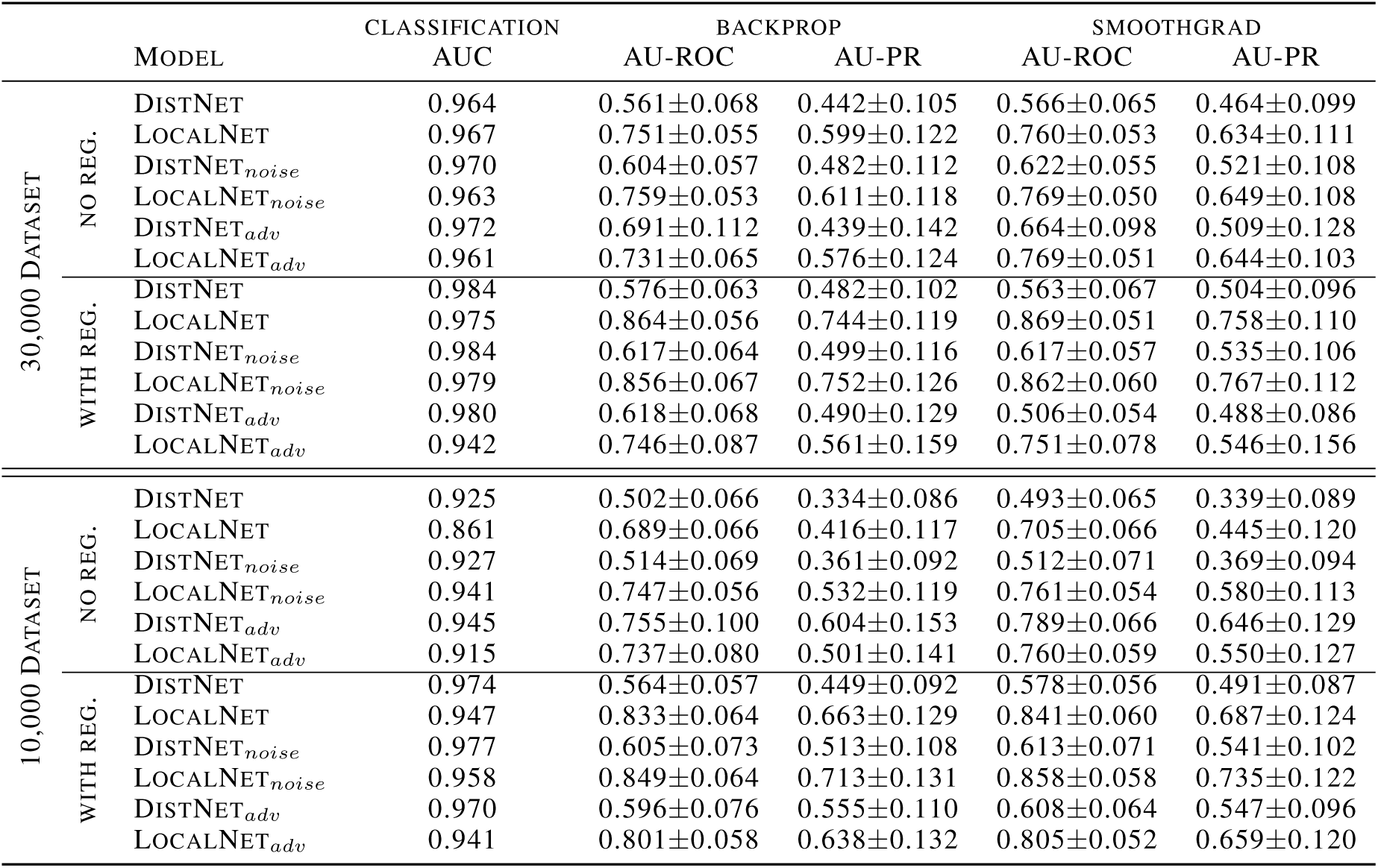
Performance summary. The table shows each model’s classification performance given by the area under the ROC curve (AUC), and the interpretability performance given by the average area under the ROC curve (AU-ROC) and the average area under the PR curve (AU-PR) using backprop and smoothgrad. Error bars are the standard deviation of the mean. Each model is annotated by training condition: standard training (no annotation), Gaussian noise injection (noise) and adversarial training (adv). Results are organized for models trained on 30,000 sequence dataset (top) and 10,000 sequence dataset (bottom), and further subdivided into whether the model is trained with or without regularization.

### Regularization significantly improves interpretability

Regularization can increase the smoothness of fitted functions in over-parameterized models, so we suspect it could improve interpretability. It has been found that regularization plays a minor role in generalization performance for neural networks (Zhang et al., 2016). We noticed a similar trend for both networks trained with and without regularization, which includes batch normalization, dropout, and *L*2-regularization (Table 1). As expected, we found that regularization significantly improves interpretability of both networks, especially for LocalNet. To see how well these findings hold for smaller datasets, we downsampled the 30,000 sequence dataset to 10,000 sequences and reran the same experiments. We find similar trends, albeit with slightly lower interpretability results (Table 1).

### Gaussian noise injection improves interpretability

Gaussian noise injection to the inputs has been found to improve the robustness of DNNs (Fawzi et al., 2016). Here, we add noise – by sampling a Gaussian distribution 𝒩 (0, 0.1) – to every nt in each sequence. A new set of noise is added at each training epoch, but not during testing. For the larger dataset, noise injection only improves DistNet on a consistent basis, while yielding mixed results for LocalNet (Table 1). For the smaller dataset, we find that noise injection during training significantly improves interpretability for each model. The most interpretable models use a combination of regularization, Gaussian noise injection and smoothgrad for both LocalNet and DistNet.

### Adversarial training has potential to improve interpretability

Another technique to improve the robustness of DNNs is adversarial training. To implement this, we first train the model for 20 epochs on clean data. Then we apply mixed adversarial training for 80 epochs, where half of each batch is clean and the other half is adversarially perturbed. We create the adversarial examples at each epoch using projected gradient descent (Madry et al., 2017) for 20 iterations, initialized with learning rate of 0.01. The maximum allowed perturbation is *ϵ* = 0.2 for *ℓ*_*∞*_ norm. In contrast to an image classification problem where adversarial examples are generated by using the least likely label as the target for the direction of the gradient, we maximize the loss for the true label to create the adversarial attack, effectively generating perturbed data that is misclassified.

In general, we find adversarial training consistently improves DistNet’s interpretability, while LocalNet exhibits mixed results (Table 1). Surprisingly, the largest gain in interpretability was for DistNet trained without regularization for the small dataset, which yields an AU-ROC and AU-PR of 0.789± 0.066 and 0.646±0.129 with smoothgrad. We verified this anomaly across multiple independent trials with different initializations (data not shown), leading us to believe that we may have unintentionally chosen a favorable combination of hyperparameters. Nevertheless, LocalNet trained with regularization and noise still yields an overall higher interpretability with an AU-ROC and an AU-PR of 0.858 ± 0.058 and 0.735± 0.122, respectively. It is challenging to find optimal hyperparameter settings for adversarial training and to determine an optimal stopping point during training. We did not fully explore alternative adversarial techniques or optimize the CNN design here. We hypothesize that further optimization could improve performance. The scope of this analysis was to explore whether adversarial training can improve interpretability.

## Conclusion

Although attribution methods have been shown to provide access to representations learned by a DNN, we raise the important issue that their interpretability is not necessarily reliable across architectures even when the DNN yields high classification performance. We showed regularization, Gaussian noise injection, and adversarial training – all of which have been demonstrated to improve robustness of DNNs in computer vision – are promising avenues to improve interpretability for genomics. Further work is required to optimize each of these training procedures specifically for genomic sequence data. Moreover, it would also be interesting to explore how other, non-gradient-based interpretability methods, such as *in silico* mutagenesis, are affected by network depth and training procedure. Further work is required to understand how to design DNNs to balance the expressiveness to fit data and the ability to also interpret them.

